# CELL-TO-CELL MODELING OF THE INTERFACE BETWEEN ATRIAL AND SINOATRIAL ANISOTROPIC HETEROGENEOUS NETS

**DOI:** 10.1101/082529

**Authors:** Gabriel López, Norma P. Castellanos, Rafael Godínez

## Abstract

The transition between Sinoatrial cells and Atrial cells in the Heart is not fully undefirst ood. Here we focus in cell-to-cell mathematical models involving typical Sinoatrial cells and Atrial cells connected with experimentally observed conductance values exclusively. We are interested mainly in the geometry of the microstructure of the conduction paths in the Sinoatrial Node. We show with some models that appropriate source-sink relationships between Atrial and Sinoatrial cells may occur according to certain geometric arrangements.

## 1. INTER-PHASE BETWEEN ATRIAL AND SINOATRIAL CELLS

It has been observed in different species that going from the center of the sinoatrial node (SAN) in the heart toward the atrium, there is a transitional zone of cells having morphological and electrophysiological properties in-between to that of typical sinoatrial (SA) and atrial (A) cells [16]. The transitional cells have an aspect intermediate between that of typical nodal cells and that of the common atrial cells. Typical nodal cells have poor development of the contractile system and is assumed in general that they do not contract, moreover the existence of connexin43 is undetectable in the SA node center (see [3] and references therein), but posses automaticity in ring its action potential. On the contrary, atrial cells do contract themselves, but they require of an stimulus in order to contract, and they contain mainly connexin-43. This characteristics are included in the cells models used in this paper. A whole range of intermediate cells have been reported, but more important to the models in this paper, cells with one end connected to SA cells and the other end with A cells have been found [16]. The basic structure conforming the cytoarchitecture of this groups of cells consists of interdigitations of nodal and atrial bundles forming histological connections between nodal and atrial myocytes at regular distances [18].

In [29] the authors, introduce a model of strands of atrial cells penetrating the SA node observed in the Pig. The model was constructed with 101 × 101 atrial and SA cells modeled with Oxsoft HEART V4.5. The lattice so constructed has a center of SA cells forming a circle of 30 cells of radius with twelve atrial interdigitations positioned at 30 deg intervals, where interdigitations are defined as sets of atrial cells at least ten cells distant from the node centre, and which are subtended by an angle of 15 deg. In that paper SA to SA conductance is g*_SA_* = 10 nS and the A to A conductance g*_A_*, varies from 10 nS to 250 nS.

In [11] the authors modeled the inter-phase in a series of 200 cells, the first 50 SA, where there is a gradient in the conductance of the Ionic currents inside the cells and also a gradient in the the membrane capacitance of SA Kurata et al. cell model (see references within [11]). In that model there is also a gradient in the conductance between SA cells varying from g*_SA_*= 20 nS to 3999.8 nS, this last value, as the authors mention, is not actually observed experimentally. For the Atrial cells they used the model of Lindblad et al. [14] with g*_A_* = 4000 nS (neither observed experimentally). In that 1D model the inter-phase is attained by one SA cell (the 50th cell in a line) connected to three branches of 50 A cells each. Then the inter-phase is achieved connecting the 50th SA cell with the 49th SA cell throughout g*_S_A* = 3999.8 nS, and connecting the 50th SA cell with three A cells with g*_SA-A_* = 4000 nS. The conductance between A cells is kept constant g*_A_* = 4000 nS. This kind of "wiggly jiggly" with parameters existing in literature over the years is not allowed in our study. Besides, note that this cell organization is not natural because one cell could fail, and in that case the entire net would fail. In the present paper g*_SA-SA_* takes values between.6 nS and 20 nS [27], and g*_A-A_* between 30 nS and 600 nS [28]. Is worth to remark that in our simulations of connections between SA and A cells conduction occurs with g*_SA-A_* as low as.6 nS.

With respect to the action potential model’s shapes, years ago, Joyner and van Capelle [13] noticed that the presence of electrical coupling among cells create transitional action potential shapes in cells near the border zone between two distinct different cell types and that the electrotonic influences make it very di cult to prove that cells in a particular region are truly transitional in terms of their intrinsic membrane properties. Accordingly with this, in our models, action potential shapes behave in some kind of transitional manner which may provide some coincidence with the behavior of transitional cells observed in vivo. We achieve this "transitional cells" through the cytoarchitecture in the model keeping the conductance as mentioned above, i.e., without any gradient between cells.

More recently in [5], Csepe et al. study the functional-structural connection between the SAN and the atria. Their studies suggest that the microstructure of the connection paths between A and SA cells plays a crucial role in human SAN conduction and contributes to normal SAN pacemaking. Our paper may be consider as a local approach to the complexity of the specialized branching myofiber tracts comprising the SA connection paths described by Csepe et al.

This paper is dived as follows. first we introduce the cell-to-cell mathematical approach that we are going to use throughout the paper; secondly we describe the models used in this paper for SA and A cells; thirdly we present the net model proposed based on observed interdigitations; fourthly we present results which show synchronization among SA and A cells with experimentally obtained conductances. Finally we establish some conclusions of our study.

## 2. MATHEMATICAL APPROACH

The cell-to-cell approach requires: a) individual cell dynamics, modeled by Hodgking-Huxley type equations. In this paper for SA node cells, we use the model of Severi et al. [23], described in section 3, and Lugo et al. model [15] for atrial cells; b) SA cytoarchitecture, in this paper we assume that the SA cells ran parallel and meet mostly end to end [24], and that each cell is connected via intercalated disks with approximately 9.1±2.2 other cells [10]. A model of idealized two-dimensional arranges of cells using a similar structure can be found in [25]; c) In order to implement the solvers an approximate number of cells in SA node is required. This number may be estimated to be in the order of millions. In the present study our approach to the transitional zone is local and the number of cells considered, varies between 30 and 78, which is enough to approximate some source-sink relations between A and SA cells in this specific part of the SAN.

Concerning to the cytoarchitecture of SA node is important not only the number of cells to which each cell is connected (already cited) but the geometric distribution of each connection. Here is important to remark that due to the use of connections matrices for the models of this paper, the inherent three dimensionality of the cytoarchitecture (see [5] where the necessity of the 3D approach in order to undefirst and the human SA node structure is amply discussed) does not require an special treatment as is the case for PDE, in which a tensor is required to describe the complex geometry distribution of cells in the heart [7], [8], [9], [21].

The equations used in this paper are of the form

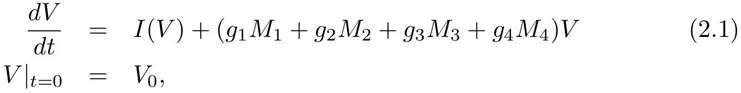

where the transposed vector 
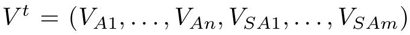
 corresponds to the Voltage of the A (*A*1 to *An*) and SA (*SA*1 to *SAm*) cells, with *n, m* values varying in different models, 
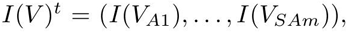
 is a vector representing the currents of A and SA cells, *M*1 is a connection matrix between A cells, *M*2 is a connection matrix between A and SA cells, *M*3 gives the connections between SA and A cells, and *M*_4_ is the connection matrix between SA cells; the constants 
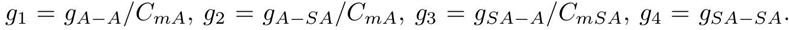
 Here *CmSA* =.000032 µF; *CmA* =.00005 µF, and *g_SA-A_*, *g_A-SA_*, *g_A-A_* are the conductance values between corresponding cells which values are specified in each model. The vector 
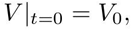
 i. e. *V* at time *t* = 0, takes values which vary randomly with normal distributions, accordingly: for Atrial cells, mean -74.2525 mV, and for SA cells, mean -58mV, with standard variation .1 mV in both cells types. In three dimensional models we keep 
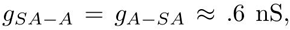
 with the precise value given in the corresponding model. We explore in section 3.2 the possibility of 
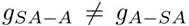
 in models of series, which can not be excluded a priori due to the possibility of existence of different kind of connexin types in the cell-to-cell connections through the transitional zone.

## 3. ATRIAL AND SINOATRIAL CELL MODELS

There are many Atrial cell models in the literature, among them: Courtemanche et al. (CRN), [4] (Human); (RNC), [22] (Canine); Nygren et al. (N), [17] (Human); Lindblad et al. (L), [14] (Rabbit), and a more recent model of Lugo et al. (LC), [15] (Human). A comparison between the models CRN, RNC, and N can be found in [6]. In this paper we use the LC model which is a refinement of N model and, basically, modifies the dynamics of the RyR2 and CA++ release of the sarcoplasmic reticulum (SR) in order to model the effect of SR release refractoriness in appearance of electromechanical alternants. The N model reconstructs action potential data that are representative of recordings from human cells. In this model the sustained outward K^+^ current determines the duration of the AP. On the other hand, the AP shape during the peak and plateau phases is determined primary by transient outward K^+^ current, I*_sus_*, and L-type Ca^2^ current.

For SA cells, in our net models, we use the Severi et al. model (S), [23] (Rabbit). This model based on up-to-date experimental data, generate Action Potential (AP) waveforms typical of rabbit SAN cells whose parameters fall within experimental ranges: 352 ms cycle length, 89 mV AP amplitude, -58 mV maximum diastolic potential, 108 ms Action Potential Duration at its half amplitude (APD_50_), and 7.1 V/s maximum upstroke velocity; and, more interesting, describes satisfactorily experimental data concerning autonomic stimulation, funny-channel blockade and inhibition of the Ca^2+^-related system by BAPTA.

### 3.1. Series Models

In [11] the authors modeled the inter-phase in a series of 200 cells, the first 50 SA, where there is a gradient in the conductance of the Ionic currents inside the cells and also a gradient in the the membrane capacitance of SA Kurata et al. cell model (see references within [11]). In that model there is also a gradient in the conductance between SA cells varying from g*_SA_*= 20 nS to 3999.8 nS, this last value, as the authors mention, exceed by far experimental values. In Atrial cells are modeled with Lindblad et al. [14] with g*_A_* = 4000 nS (neither observed experimentally). In that 1D model the inter-phase is attained by one SA cell (the 50th cell in a line) connected to three branches of 50 A cells each. In that model the inter-phase is achieved connecting the 50th SA cell with the 49th SA cell throughout g*_SA_* = 3999.8 nS, and connecting the 50th SA cell with three A cells with g*_SA-A_* = 4000 nS. The conductance between A cells is kept constant g_A_ = 4000 nS.

In [19] Oren and Clancy, study a series model (among others models) of 60 cells. In this model a gradient in coupling, shifts the site of pace maker initiation towards the periphery of SA node, a result which, as the authors mention, is not consistent with experimental observations.

In this paper we assume that SA cells are synchronized in phase, as theoretically predicted in the well known paper of Torre [26]. Moreover for S model varying randomly *V*_0_ in each simulation (up to 20 simulations) we obtained synchronization in phase of 100 SA cells, even before the first cycle ended. For the numerical simulations we considered different geometric arrangements: in series, forming an anullus, and even with a random matrix of connections (which is histologicaly improbable). In order to model connections between 100 SA cells we take in equation (2.1) *M_1_* = *M_2_* = *M_3_* = **0**_100×100_ where **0**_100×100_ is the zero matrix of size 100×100, and *g*_4_ =.6nS/*CmSA*.

Hence in a series model (see figure 1) the transition zone of the SAN can be modeled locally with just one SA cell using the S model for SA cells. We are interested in modeling without introducing coupling parameters not observed experimentally, so we keep in our simulations, SA coupling values between.6 and 25 nS [27] (Rabbit). For this model equation (2.1) becomes simply

**Figure 1.**
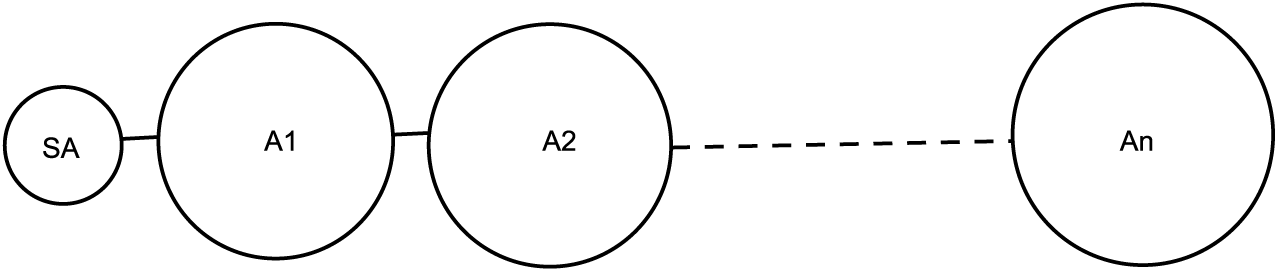
Series arrangement of cells.

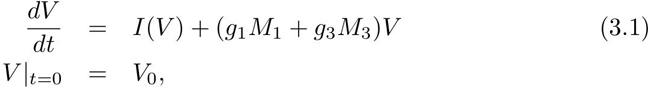

where *M*_1_ is a tridiagonal matrix with main diagonal (-2,-2,…,-2) and sub and super diagonals (1,…,1); and *M*_3_ is the matrix with first row (-1,0,…,0,1), last row (1,0,…,0,-1), and an zero matrix in between this two rows.

### 3.2. Series models analysis

The figures and a detailed analysis of this subsection are included in the Appendix. Using the N model, conduction occurs if, for instance *g_A-SA_* = *g_SA-A_* = 6 nS. We observe nevertheless, that conduction occurs if *g_A-SA_* ± *g_SA-A_*, for instance, if *g_A-SA_* is 10 fold *g_SA-A_* = 6 nS. For more examples in which conduction fails in this series model consult the Appendix. Curiously, the A cells AP profile behave as SA cells AP pro le if *g_A-SA_* = 100*g_A-SA_* = 100×.006 nS.

We call selective diffusion to the phenomenon occurring in simulations in which conduction is achieved if *g_A-SA_ ≠ _SA-A_*. The question here is if this kind of selective diffusion may actually occur in real cells, to the best of our knowledge this is an open question to date, the mechanisms of coupling mechanisms remain poorly understood. Recent studies [2] indicate that connexins as Cx43 may exert different functions besides gap junctions formation, including non-cannonical functions in the function of sodium channel. So to-date the possibility of selective diffusion can not be excluded a priori.

Another form in which conduction fails occurs when some of the cells do not reach threshold for instance in a model with .4*g_A-SA_* =.1*g_SA-A_*. (See the Appendix).

A more interesting model is obtained if we abandon the one dimensional restriction of the series, for instance, when six or seven SA cells are attached to a series of A cells as shown in figure 7. This models contradict the claim of Inada et al. [11] that suggest that a gradient in both electrical coupling and cell type is necessary for the SAN to drive the atrial muscle. The left hand side plot in figure 7 shows one of seven SA cells (the first) connected in a series with atrial cells, with same conductance *g_SAA_* = *g_AA_* =6nS. Note nevertheless, that the action potential of the first four atrial cells show a transitory cell profile, this cells are typical A cells without any gradient, so that conduction is possible and this phenomenon may explain why in vivo there appear transitional cells similar to atrial cells.

### 3.2.1. Conduction is not achieved

In the next section we will study models in more than two dimensions. As an introduction we mention that if a series is divided in order to branch, then seven SA cells are not enough to propagate the AP. Accordingly, we constructed two models in which propagation fails. In the first model an A cell is connected to seven SA cell and to two A cells which in turn are connected to two disconnected series, to illustrate this, in figure 3, A1 is connected to A2 and to A25; A2 is connected to a series from A3 to A24; and A25, in turn, is connected to a series from A26 to A50. The way in which conduction fails is shown in figure 8. In the second model in which conduction fails A1 is connected to seven SA cells and to A2 cell, then A2 branches into two disconnected series. The important issue here is that conduction occurs if we abandon the series model. In fact, in figure 9 the same bifurcate series conducts AP as long as 4 SA cells are connected to each A2 and A25 cells as in figure 2. Note that we only plot 1 SA cell and the series A2 to A14, and cells A23, A24.

**Figure 2.**
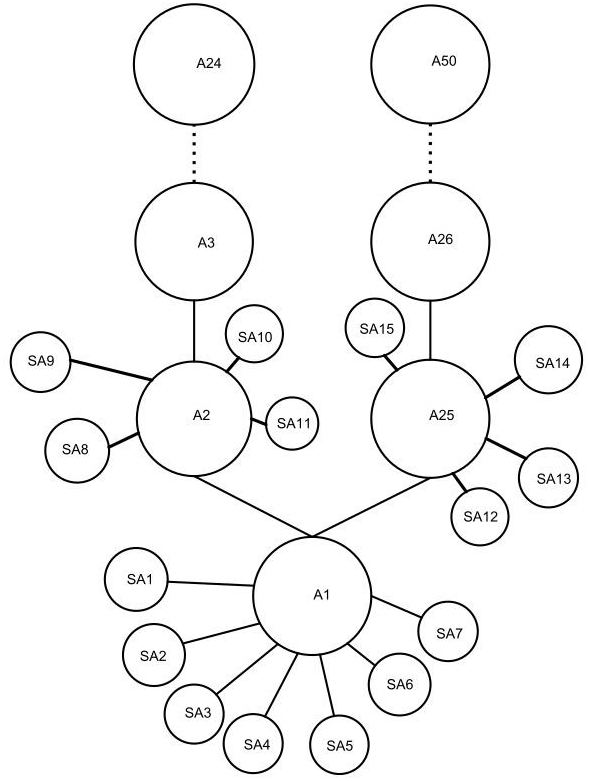
Two disconnected series in which conduction does occur.

**Figure 3.**
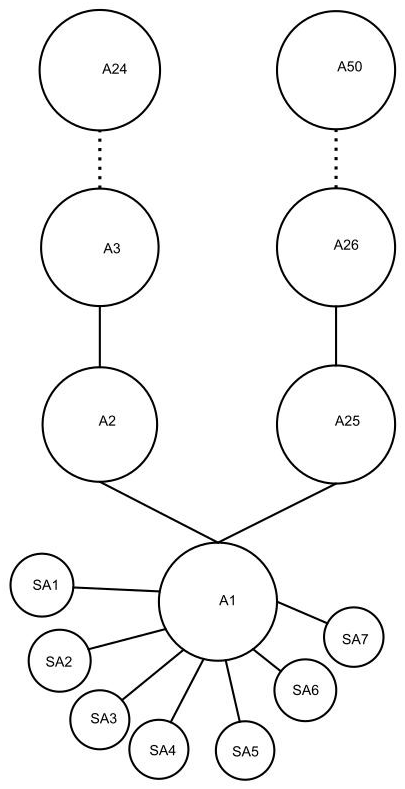
Two disconnected series in which conduction does not occur.

### 3.3. Tree Net Models

We call a Tree Net a model in which each cell is connected with two or three other cells forming branches. In Tree net models the number of cell connected spreads rapidly. For instance in the tree model in which each cell is connected with three other cells, after twenty steams we would have (3^21^-1)/2 = 5.2e+9 cells connected and activated by one single SA cell, a number much larger than the entire number of cells in the entire heart (the number of cells in a healthy human heart is about 2 billion (2e+9), [1]). Therefore, the Tree Net Model is somehow an unrealistic model for the bundle formed by SA node and Atrial cells, nevertheless, as a first approximation, it will provide some interesting information. We clarify this point, in order that an small SA node paces the heart it should be expected that A cells do ramify, but not at every cell, the manner in which Atrial cells do ramify is, to the best of our knowledge, unknown, but certainly ramifications do no occur at each cell. Nevertheless in modeling the transition between SA and A cell we found that each ramification requires a minimum number of SA cells in order to drive a tree of A cells if we want to keep *gSA_A_* within experimental values and even, if we want to keep this parameter value constant.

In figure 5, a tree like that in figure 4 (i. e. with ramifications in each node) is paced if A1 is connected with seven SA cells and cells A2; A3 are connected with another four SA cells each. A comment is worth here, clearly 7 cells connected with only one cell (as in figure 2) is impossible in one dimensional series models and very difficult even in two dimensional models given SA and A cells sizes, but this number of connections may occur in three dimensional models without geometrical complications, as actually they do happen. So our model is intrinsically a model in three dimensions. But more importantly our model suggest that in order to obtain conduction with *g_SA-A_* =.5 nS constant, the branch must be connected to a minimum number of SA cells which in our model is 15 at the first branch in a tree. Actually we have to mention that is possible to pace the net with 15 SA cells connected with A1 cell and this cell connected with a net of A cells, but we obtain subthreshold behavior in A cells, even with *g_SA-A_* = 0.57 nS, which clearly indicates that the geometry of the net is extremely relevant (see figure 6) for conduction.

**Figure 4.**
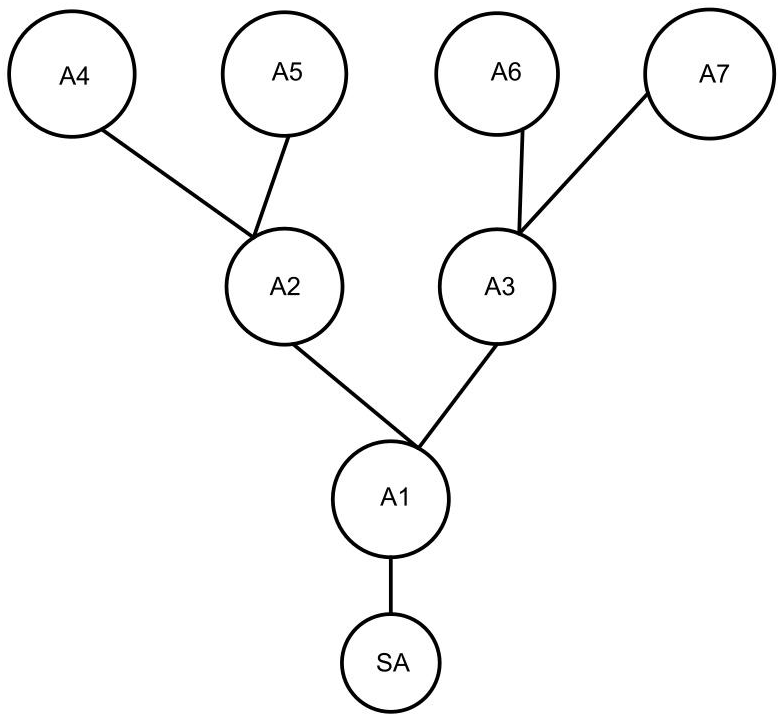
Tree with two-type ramifications

**Figure 5.**
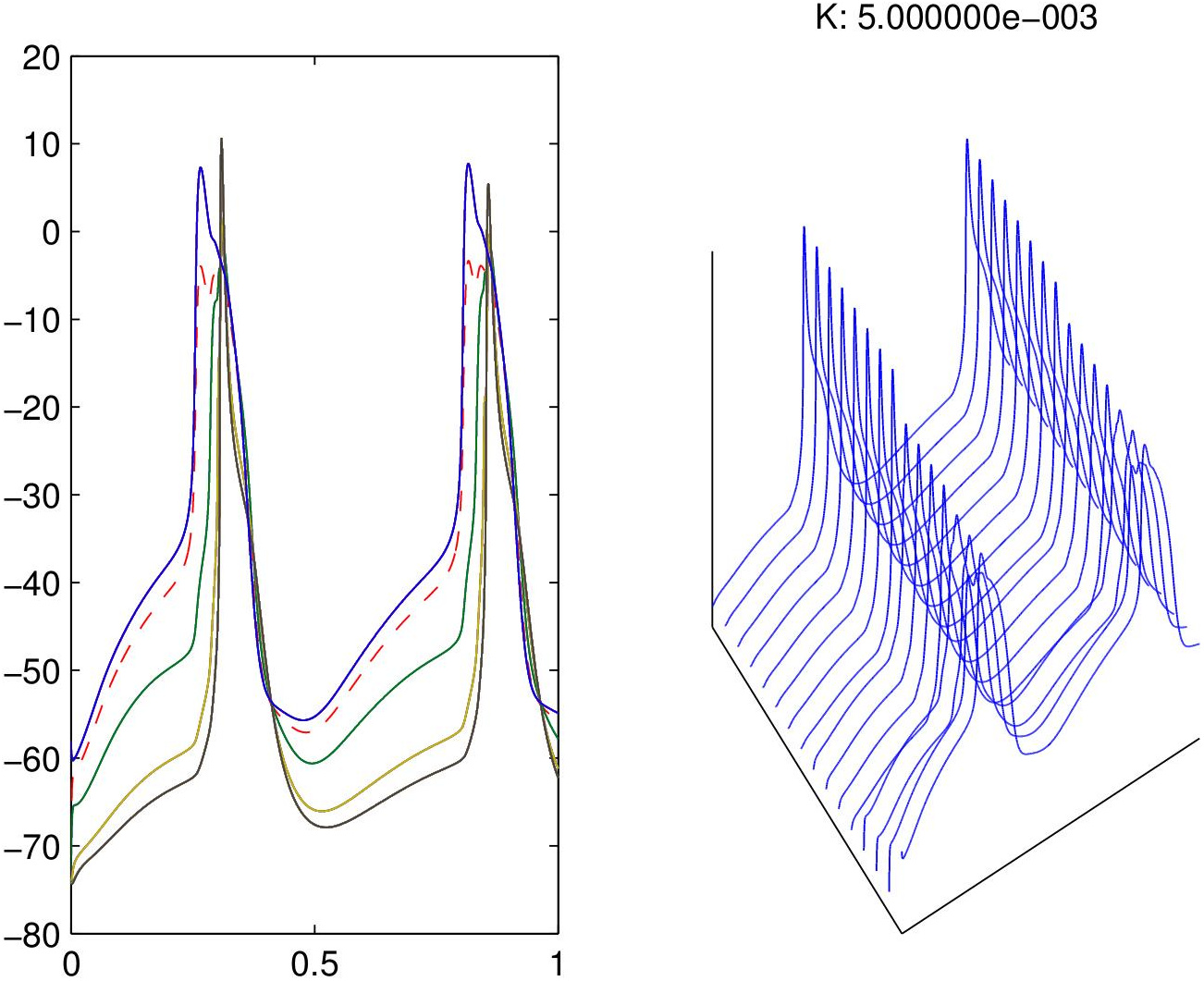
Tree with two-ramifications at each cell.

**Figure 6.**
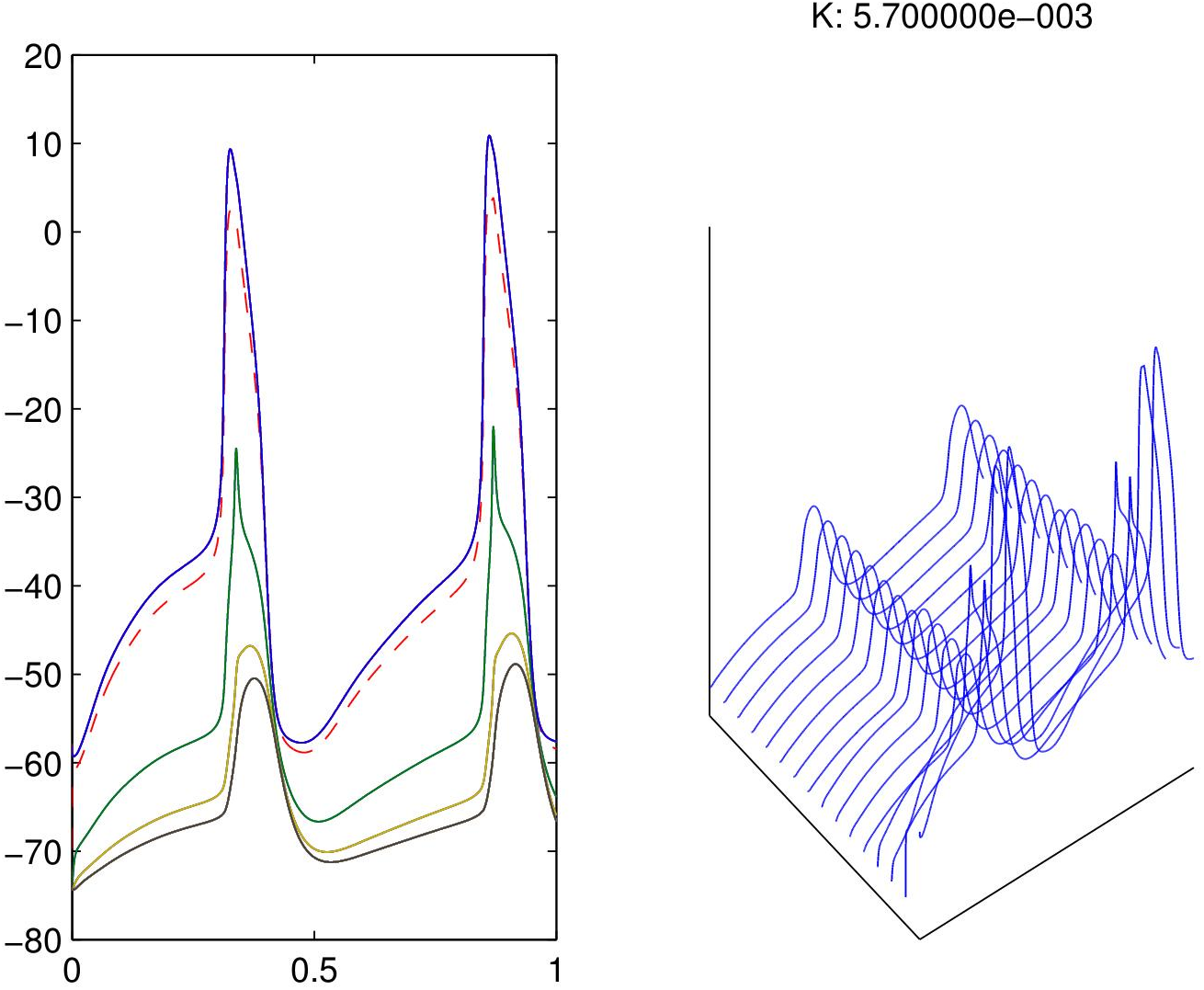
Tree with two-ramifications, subthreshold A cells

Finally, in numerical simulations we obtain that conduction is possible if we allow *g_SA-A_* ± *g_A-SA_* as in series models. In figure 10, *g_A-A_* = *g_SA-A_* =.6 nS, but g_A-SA_ = 14 nS. Notice that A2, A3 cells present two peaks which may be consider as an anomalous behavior.

Mathematically, at least qualitatively, the behavior of the nets of this section can be described by the existence of a supercritical Andronov-Hopf bifurcation [12], i. e., a bifurcation of an equilibrium state (sink, due to the A cells) passing to a small-amplitude limit attractor (like that in figure 6 due to the sources in SA cells), and as the number of SA cells connected increases, the amplitude of the limit cycle increases to become a full size limit cycle of each cell in the net (like those in figure 6 and figure 7). Remarkably the bifurcation parameters give also the structure of the tree in each model, structure which coincides, at least locally, with interdigitations of A and SA cells.

**Figure 7.**
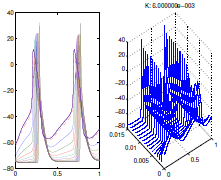
Model in which conduction occurs.

**Figure 8.**
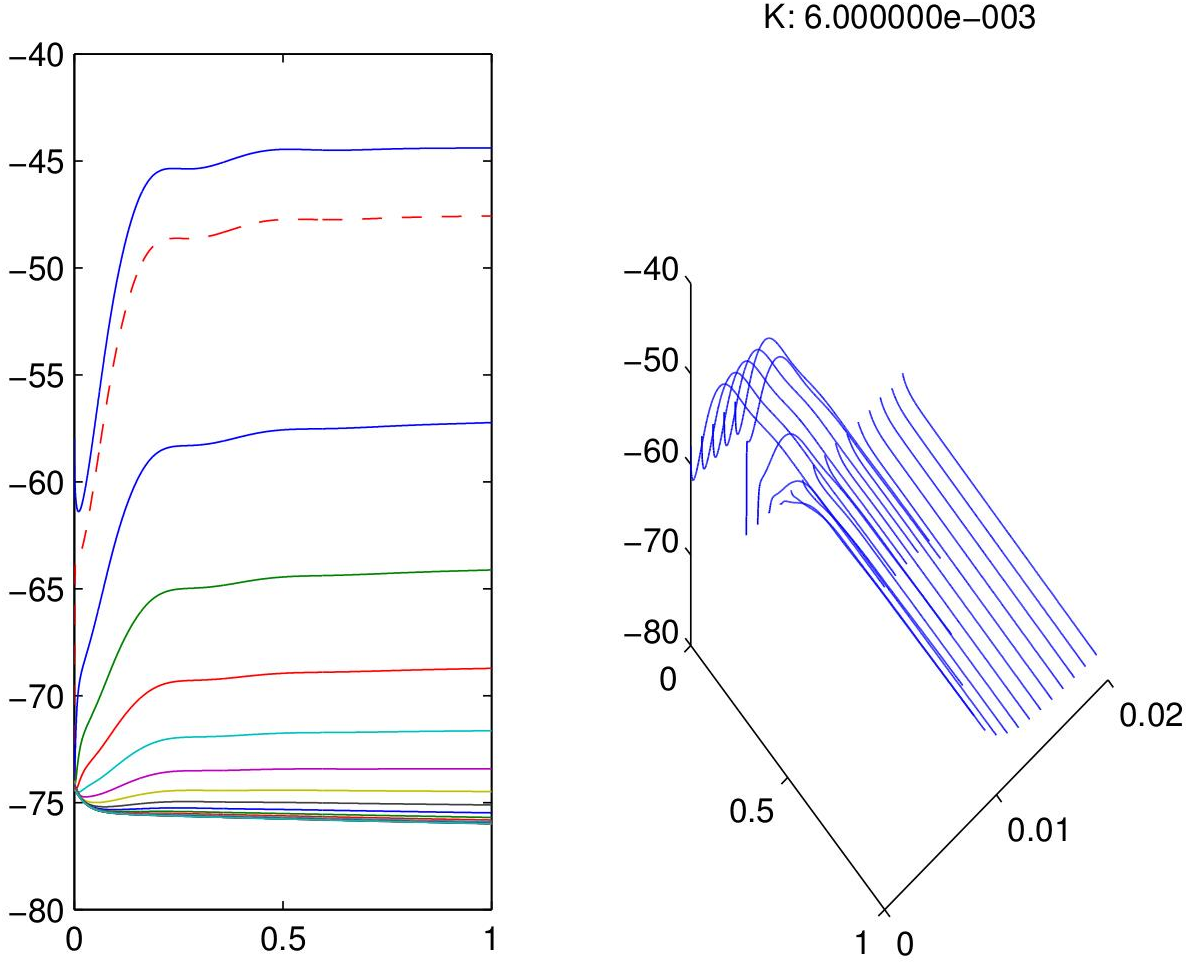
Series model in which conduction fails.

**Figure 9.**
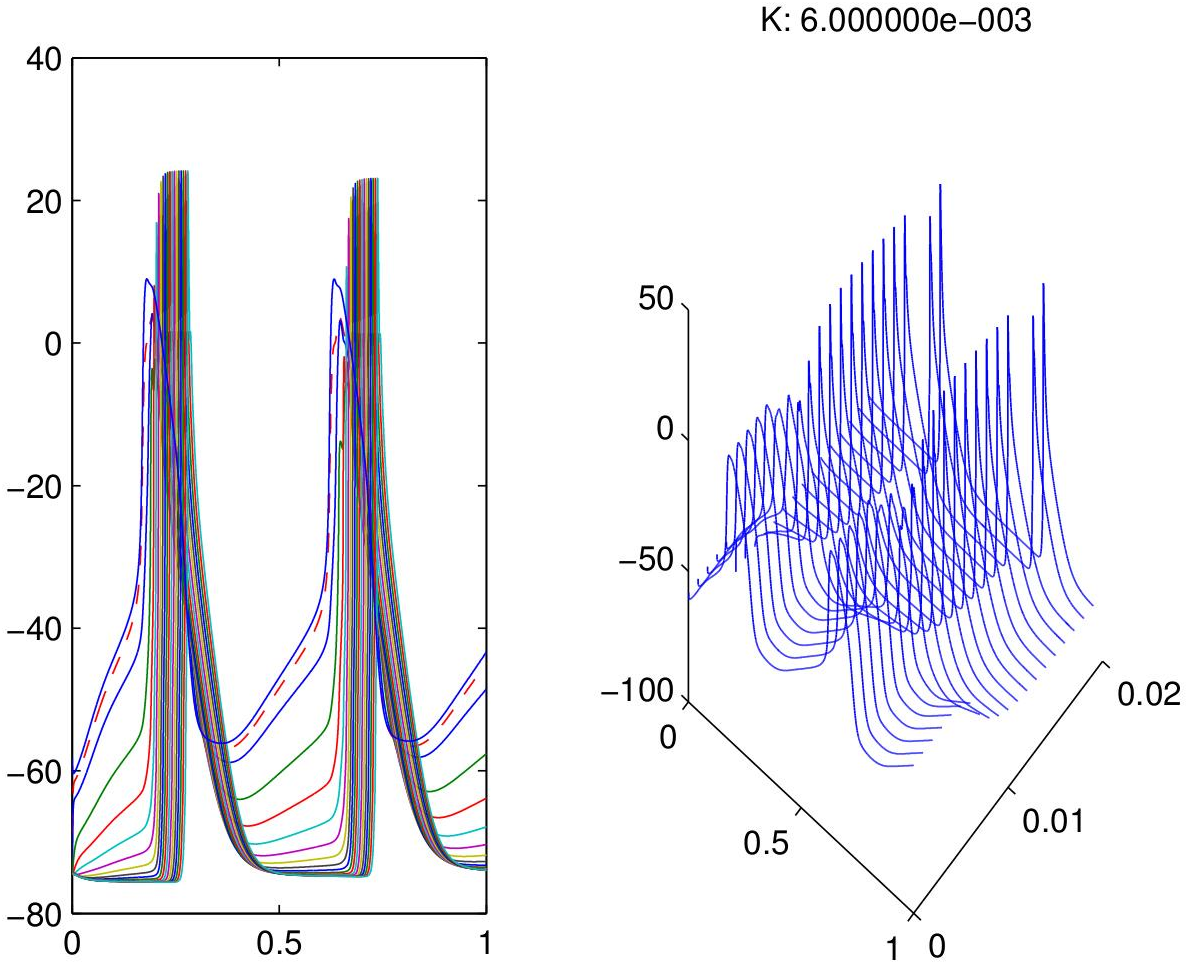
A bifurcated series which conducts AP

**Figure 10.**
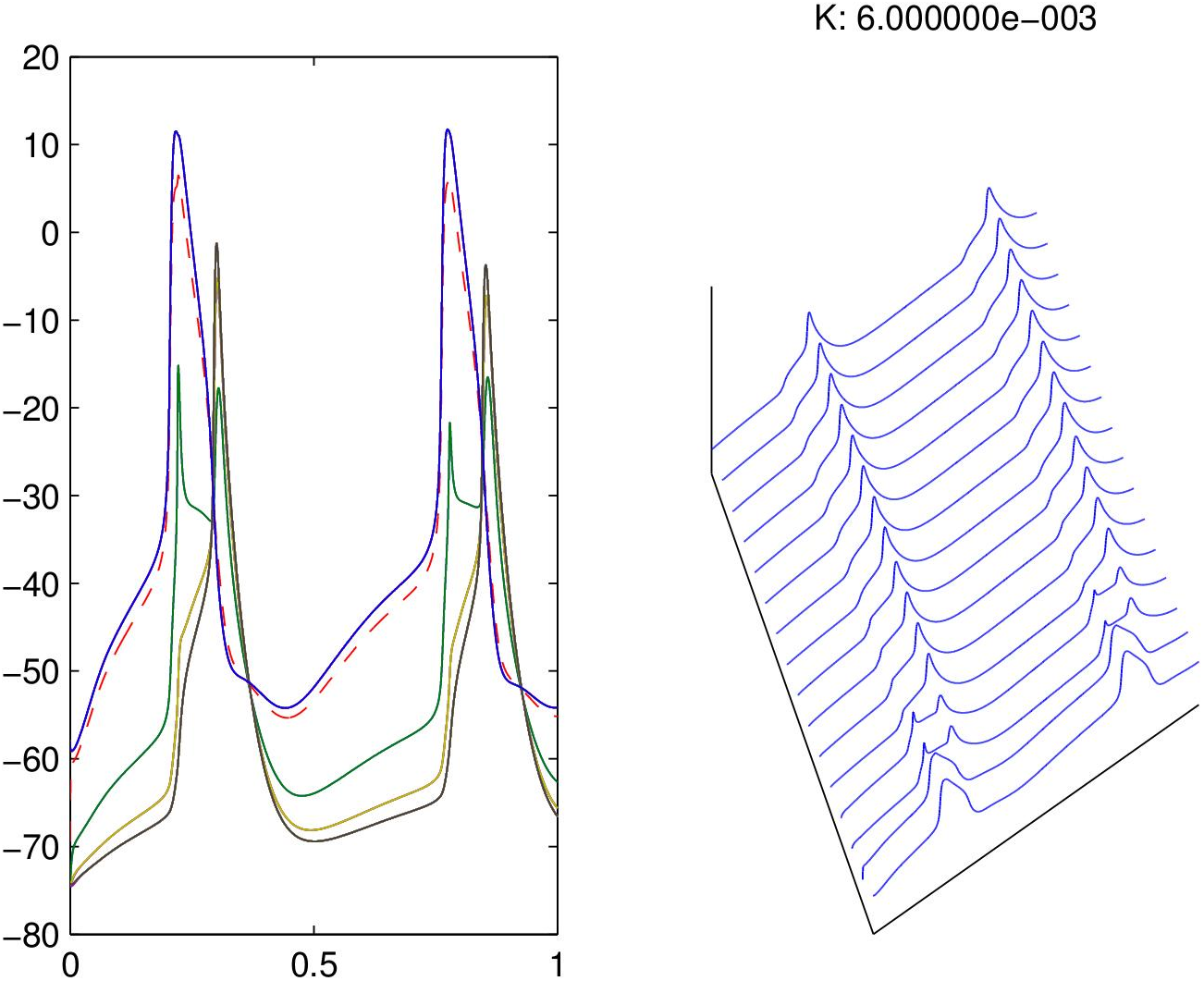
Tree with *g_SA-A_* ± *g_A-SA_*.

## CONCLUSIONS

In the literature some models of the transitional zone between SA and A cells require non experimental conductance values in order to obtain propagation of the AP generated in the SA node. On the contrary, in this paper we obtained models which do not require artificial parameter values to achieve propagation. Our models mimic locally the strand structure (interdigitations) observed and studied in classical papers as [29], and in recent papers as [5]. On the other hand, our models are compatible with mosaic models but may allow the implementation of gradients in some variables as long as this variables do no modify drastically observed conductance values. Mathematically, our successfully conducting AP models may be explained by a supercritical Andronov-Hopf bifurcation associated to a correct source-sink relationship in the transitional zone. Remarkably, the bifurcation parameters of the Andronov-Hopf bifurcation give a geometrical distribution between A and SA cells which is compatible with interdigitating bundles structure of the transitional zone. Hence we conclude that in modeling the cytoarchitecture of the transitional zone between SAN and Atrial zone is essential for smooth propagation of AP to include geometrical arrangements, accordingly with some bifurcation parameters of the dynamical systems theory.

## REFERENCES

[1] Adler CP, Costabel U. Cell number in human heart in atrophy, hypertrophy, and under the influence of cytostatics. Recent Adv Stud Cardiac Struct Metab. 1975;6:343–55.

[2] Agullo-Pascual E., Delmar M., The “non-canonical” functions of Cx43 in the heart. J Membr Biol. 2012 August; 245(8): 477–482. Vol 47(7): 907–918, 1999.

[3] Coppen S. R., Kodama I., et al Connexin 45, a Major Connexin of the Rabbit Sinoatrial Node, Is Co-expresed with Connexin43 in a restricted Zone at the nodal-Crista Terminalis Border. The Journal of Histochemistry & Cytochemestry Vol 47(7):907–918, 1999.

[4] Courtemanche M., Ramirez R., Nattel S., Tonic mechanisms underlying human atrial action potential properties: insights from mathematical model. the American Physiological Society, H301–H321 (1998).

[5] Csepe T. A., Zhao J., et al. Human sinoatrial node structure: 3D microanatomy of sinoatrial conduction pathways. Progress in Biophysics and Molecular Biology 120(2016) 164–178.

[6] Cherry E. M., Hatings, H. M., Evans S. T., Dynamics of human atrial cell models: Restitution memory, and intracellular calcium dynamics in single cells. Progress in Byophysics and Molecular Biology 98 (2008) 24–37.

[7] Colli Franzone P., Pavarino L., A parallel solver for reaction-diffusion systems in computational electrocardiology. Mathematical Models and methods in Applied Sciences Vol. 14, No. 6 (2004) 883–911.

[8] Colli Franzone P., Guerri L., Spreading of Excitation in 3-D Models of the Anisotropic Cardiac Tissue. I. Validation of the Eikonal Model. Mathematical Biosciencies 113:145–209 (1993).

[9] Colli Franzone P., Guerri L., et al., Spreading of Excitation in 3-D Models of the Anisotropic Cardiac Tissue. II. Effects of Fiber Architecture and Ventricular Geometry. Mathematical Biosciencies 147:131–171 (1998).

[10] Hoyt R. H., Cohen M.L., et al. Distribution and Trhee-Dimensional Structure of intercellular Junctions in Canine Myocardium. Circ Res. 1989; 64: 563–574.

[11] Inada, S., Zhang, H. et al. Importance of Gradient in Membrane Properties and Electrical Coupling in Sinoatrial Node Pacing. PLOS ONE, April 2014 Vol 9 Issue 4 e94565.

[12] Izhikevich, E. M., Dynamical Systems in Neuroscience, the geometry of excitability and Bursting. Massachusetts Institute of Technology 2010.

[13] Joyner R. W., van Capelle F. J. L., Cells Propagation Through electrically Coupled How a Small SA Node Drives a Large Atrium. Biophys. J. Vol 50 Dec 1986 1157–1164.

[14] Lindblad D. S., Murphey C. R., Clark J. W., Giles WR., A model of the action potential and underlying membrane currents in a rabbit atrial cell. Am J Physiol. 1996 Oct;271(4 Pt 2):H1666–96.

[15] Lugo C. A., Cantalapiedra I. R., Pearanda A., Hove-Madsen L., Echebarria B., Are SR, Ca content uctuations or SR refractoriness the key to atrial cardiac alternans?: insights from a human atrial model. Am J Physiol Heart Circ Physiol. 2014 Jun 1;306(11):H1540–52. doi:10.1152/ajpheart.00515.2013. Epub 2014 Mar 7.

[16] Masson-Pévet, M. A., Bleeker, W. K., et al. Pacemaker Cell Types in the Rabbit Sinus Node: A correlative Ultrastructural and Electrophysiological Study. J Mol Cell Cardiol 16, 53–63 (1984).

[17] Nygren A., Fiset C., Firek L., Clark J. W., Lindblad D. S., Clark R. B., Giles W. R., Mathematical Model of an adult Human Atrial Cell. The Role of K^+^ Currents in Repolarization. Circ Res. 1998; 63–81.

[18] Oosthoek, P. W., Virágh, S. et al. Immunohistochemical Delineation of the Conduction System I: The Sinoatrial Node. Circ Res 1993;73:473–481.

[19] Oren, R. V., Clancy, C. E. Determinants of Heterogeneyty, Excitation and Conduction in the Siniatrial Node: A model Study. PLOS Computational Biology, December 2010, Vol 6 Issue 12 e1001041.

[20] Orthof T., The mammalian sinoatrial node. Cardiovascular Drugs and Therapy 1: 573–597, 1988.

[21] Pan lov A., Keener J., Re-entry in three-dimensional Fitzhugh-Nagumo medium with rotational anisotropy. Physica D 84 (1995) 545–552.

[22] Ramirez R. J., Nattel S., Courtemanche M., Mathematical analysis of canine atrial action potentials: rate, regional factors, and electrical remodeling. Am J Heart Circ Physiol 279: H1767–H1785, 2000.

[23] Severi S., Fantini M., Charawi L. A., DiFrancesco D. An updated computational model of rabbit sinoatrial action potential to investigate the mechanisms of heart rate modulation J Physiol 590.18 (2012) pp 44834499–4483.

[24] Shimada T., Kawazato H., et al. Cytoarchitecture and Intercalated Discs of the working Myocardium and the Conduction System in the Mamalian Heart. The anatomical Record Part A 280A:940–951 (2004).

[25] Spach M., Heidlage J. F., The Stochastic Nature of Cardiac Propagation at a Microscopic Level. Circ Res., 76(3): 366–380, Mar 1995.

[26] Torre, V. A theory of Syncchronization of Heart Pace-maker Cells. J. theor. Biol. (1976), 55-71. 1998; 97: 1623–1631 doi: 10.1161/01.CIR.97.16.1623

[27] Verheule, S., van Kempen M. J., Postma, S. et al. Gap Junctions in the rabbit sinoatreial node. Am J Physiol Heart Circ Physiol (2001) 280; H2103–2115.

[28] Verheule, S., van Kempen M. J. et al. Characterization of gap junction channels in adult rabbit atrial and ventricular miocardium. Circ Res 1997(80); 673–681.

[29] Winslow, R. L., Jongsma, H. J., Role of tissue geometry and spatial localization of gap junctions in generation of the pacemaker potential. Journal of Physiology (1995) 487. P.

